# The genomic epidemiology of SARS-CoV-2 in Palestine

**DOI:** 10.1101/2020.10.26.355677

**Authors:** Nouar Qutob, Zaidoun Salah, Damien Richard, Hisham Darwish, Husam Sallam, Issa Shtayeh, Osama Najjar, Mahmoud Ruzayqat, Dana Najjar, François Balloux, Lucy van Dorp

## Abstract

Severe acute respiratory syndrome coronavirus 2 (SARS-CoV-2), the novel coronavirus responsible for the COVID-19 pandemic, continues to cause significant public health burden and disruption globally. Genomic epidemiology approaches point to most countries in the world having experienced many independent introductions of SARS-CoV-2 during the early stages of the pandemic. However, this situation may change with local lockdown policies and restrictions on travel leading to the emergence of more geographically structured viral populations and lineages transmitting locally. Here, we report the first SARS-CoV-2 genomes from Palestine sampled from early March, when the first cases were observed, through to August of 2020. SARS-CoV-2 genomes from Palestine fall across the diversity of the global phylogeny, consistent with at least nine independent introductions into the region. We identify one locally predominant lineage in circulation represented by 50 Palestinian SARS-CoV-2, grouping with isolated viral samples from patients in Israel and the UK. We estimate the age of introduction of this lineage to 05/02/2020 (16/01/2020 - 19/02/2020), suggesting SARS-CoV-2 was already in circulation in Palestine predating its first detection in Bethlehem in early March. Our work highlights the value of ongoing genomic surveillance and monitoring to reconstruct the epidemiology of COVID-19 at both local and global scales.

## Introduction

Severe acute respiratory coronavirus 2 (SARS-CoV-2), the novel coronavirus responsible for the Coronavirus disease 2019 (COVID-19) pandemic has spread rapidly around the world since its emergence towards the end of 2019 in China^1–3^. Since then over 100,000 SARS-CoV-2 genome assemblies have been made available thanks to the global efforts of public health agencies and research teams^4,5^. This large and growing resource has brought genomics to the forefront as a method to understand both the ongoing evolution of the virus but also as a surveillance and epidemiological tool^6^.

Genomic data can be a rich source of information to inform on a variety of key epidemiological parameters such as the age and geographic origins of epidemics, their relative growth rates, to distinguish persistent infections from reinfections and to inform on the relative contributions of imported cases compared to sustained community or cryptic transmission. A wealth of genomic studies of SARS-CoV-2 from the more local^3,7–16^ through to continental^17,18^ and global scales^1,19^ have consistently pointed to most densely sequenced countries around the world having experienced a number of independent introductions, seeding local transmission chains which are subsequently maintained or may go extinct. For example, analyses of a large UK cohort identified a minimum of 1,356 independent introductions of SARS-CoV-2 into the UK, with first detected transmission events peaking in mid to late March 2020^10^. Analyses of the early Washington State outbreak could identify, using a spatial Bayesian framework, introductions from Hubei province, China, in late January to early February; with similarly early outbreaks in Northern Italy likely deriving from introductions from China over a comparable time period^17^.

With increasing use of non-pharmaceutical interventions to tackle COVID-19, including travel bans, social distancing measures and local/nationwide lockdowns, the nature of SARS-CoV-2 transmission may be altered from that reconstructed early in the pandemic. In particular, more recent genomic epidemiology studies have identified the presence of closely related sets of viruses in circulation which may define within-country spatial infection clusters, sometimes deriving from known close contact events^20^. A recent example is the reappearance of SARS-CoV-2 in New Zealand despite the virus not having been observed for 102 days prior to its re-emergence in the community. While SARS-CoV-2 samples collected in New Zealand during the ‘first wave’ derived from multiple imports, predominately from North America^16^, early analysis of samples collected during the August 2020 outbreak suggest the secondary outbreak consists largely of very closely related B.1.1.1 assigned isolates (https://nextstrain.org/ncov/oceania?c=region)^6^.

Some regions of the globe have conducted extensive genomic surveillance of SARS-CoV-2. For example, the UK viral population has been sampled to unprecedented depth (>40,900 complete assemblies on GISAID as of 19/10/2020). Conversely, the diversity of SARS-CoV-2 strains circulating in other regions of the world remains under sampled and under studied. A wider geographical coverage of SARS-CoV-2, including genomic samples from additional countries is precious as it may in time facilitate comparisons over many nations, characterised by different climate, pandemic mitigation strategies, human population densities as well as the age/health status of the general population.

The government of Palestine declared an emergency period for one month on March 5^th^ 2020, after seven Palestinians tested positive for SARS-CoV-2 in Bethlehem on March 4^th^. A curfew was declared, quarantining the population except in cases of emergency. The state of emergency was extended for one month thrice with the last such announcement on May 4^th^. On May 25^th^, the restrictions were eased following a decline in cases and a reduction of the rate of positive tests in Palestinian workers returning from Israeli areas. Seroprevalence, as measured up to July 2020 remains low^21^.

However, cases surged again during July, with the epicentre of the epidemic in Hebron accounting for over 70 % of active cases. On the 3^rd^ of July, a ten-day complete lockdown was declared across the entire West Bank. On the 12^th^ of July, a complete lockdown for five days was declared in Hebron, Bethlehem, Ramallah and Nablus governorates. Movement between all governorates was prohibited until the 27th July, with a nighttime and weekend curfew imposed on residents except for a few permitted services. All social public gatherings and transportation between governorates were prohibited. However, after the 13^th^ of July, the government of Palestine announced an ease in the restrictions allowing small businesses to reopen, subject to restrictions, and commercial movement between governorates. A state of emergency was extended for a tenth time on October 1^st^. As of the 19^th^ of October, 58,539cases and 478 deaths have been reported by the Palestinian Ministry of Health (MOH).

To better understand the epidemiology of early introduction and transmission events of SARS-CoV-2 in Palestine, through to the spring epidemic and its aftermath until late summer, we generated high quality genomic assemblies for 69 SARS-CoV-2 isolates sampled from patients between the 4th of March through to the 19^th^ of August 2020. We phylogenetically placed these samples in the context of 54,804 global samples available on GISAID from the end of August 2020 (25/08/2020). This allowed us to quantify the minimum number of introductions of SARS-CoV-2 into Palestine and to identify a sizeable local transmission cluster sustained since its appearance, which we estimate to predate significantly the first documented COVID-19 cases in Palestine.

## Results

### Palestine SARS-CoV-2 samples fall across the diversity of global clusters

Our data comprises 69 SARS-CoV-2 genome samples spanning from the early stages of the COVID-19 epidemic in Palestine from March through until late August, collected in 17 locations (7 governorates) (**Figure S1**, **Table S1**). The average difference between any two samples was 11.6 mutations (95% CI: 8.13-18.10), though with detectable structure in SNP sharing patterns often following the governorate of sampling (**Figure S2**). A complete list of mutations carried by each isolate is provided in **Table S2**.

When placed in a large global phylogeny of SARS-CoV-2, the 69 isolates from Palestine fall into six lineages, defined by the Pangolin dynamic lineage classification tool^22^, interspersed over the global phylogenetic tree (**Figure 1**, **Table S1**). This includes five ‘singleton’ isolates that are phylogenetically unrelated to any other isolate obtained from Palestine as well as a pair of related isolates assigned to the B.1.9 lineage. The pairwise SNP differences across a random sub-sample of the global alignment (mean 12.8; 95% CI 8.2-19.4) show no significant differences to those observed in Palestine, meaning our dataset can be considered as a representative random sample of the SARS-CoV-2 genomic diversity in circulation globally.

**Figure 1.**
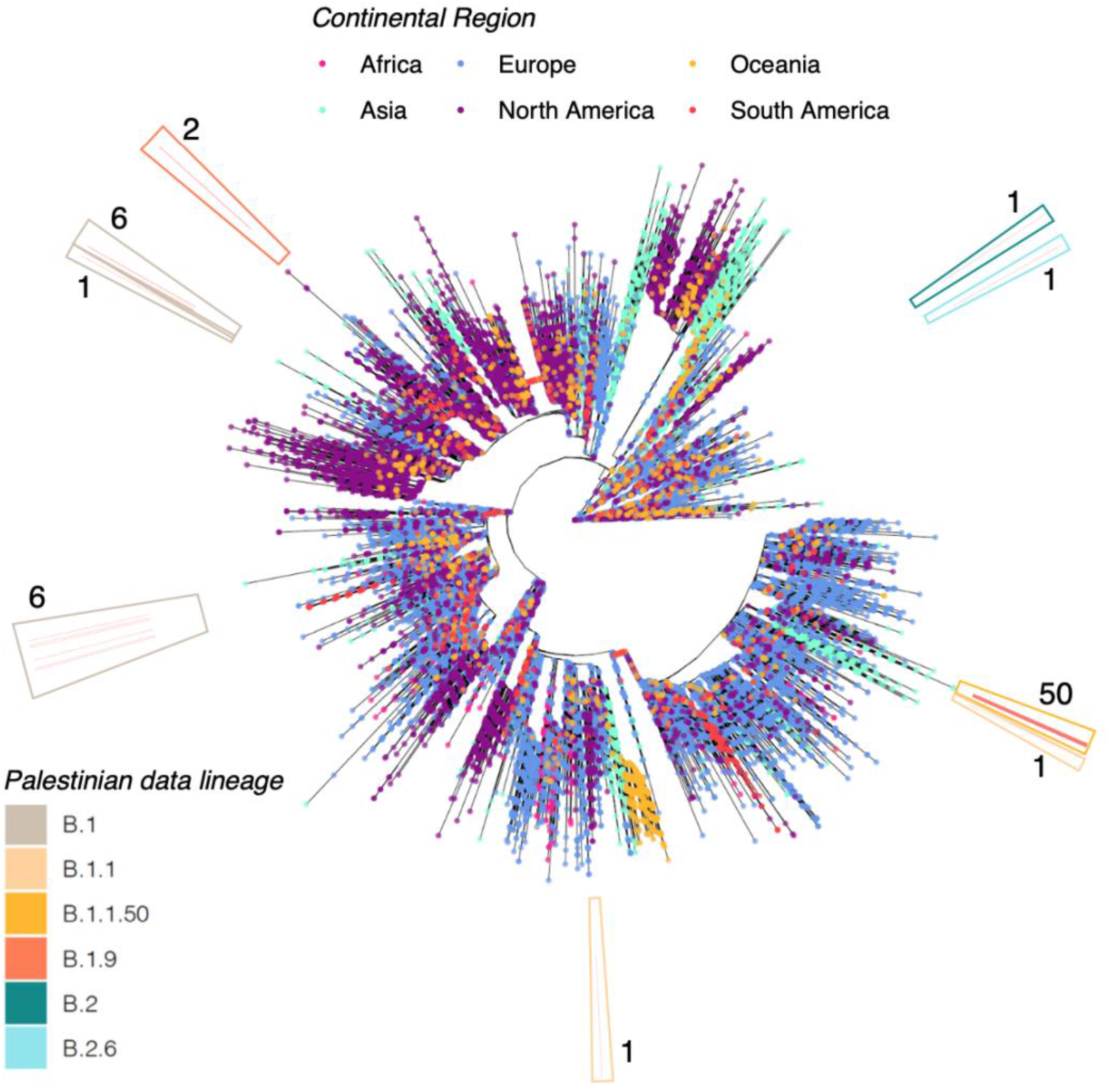
Phylogenetic placement of data collected in this study in the context of a large global phylogeny of 54,804 SARS-CoV-2 assemblies. Tip colour provides the continental region of sampling as given by the legend at top. The outer ring highlights the number of samples in different phylogenetic clades with the outer border providing the Pangolin lineage assignments as per the key at bottom left.

### From the global to the local

We additionally observe three phylogenetic clades comprising multiple closely related SARS-CoV-2 strains sampled in Palestine. This includes two distinct B.1 associated lineages of six strains and one large cluster of B.1.1.50 SARS-CoV-2 (**Figure 1**). The first B.1 associated clade comprises five samples from Bethlehem and one from Ramallah spanning from the 4^th^ of March through to the 29^th^ of March. Three samples are zero SNPs apart (sample identifiers: 28, 16, 19) with two collected in Bethlehem, both on the 16^th^ of March, and one from Ramallah 13 days later (**Figure S3**). These three samples fall within a cluster of 110 SARS-CoV-2 which are genetically strictly identical despite having been sampled in 21 different countries between the 4^th^ of March to 26^th^ of April (**Figure S4**). Within this cluster we identify that samples 16 and 28 share 29 minority variants (among which two are found only in those two strains) which might be indicative of local transmission. The second B.1 associated clade of six samples includes two SARS-CoV-2 sampled from Tulkarem on the 4^th^ of March (genetically identical), two samples from Bethlehem and two from Jerusalem all sampled on the 31^st^ of March (**Figure S5**).

### B.1.1.50 transmission cluster

The majority of our samples (n=50; 73%) however fell into a single, tight clade of closely related strains also including one sample from the UK and eight from Israel (**Figure S6**) spanning a collection period from March to August 2020 (**Figure S7**) and encompassing an average of 8.5 (95% CI 5.8-14.6) pairwise SNPs (**Figure S8**). While we do not detect significant accumulation of mutations over the sampling period in this clade, both the global phylogeny and a subset sample of 1,220 B.1.1 isolates including this clade exhibit highly significant temporal signal following randomisations of sampling date (p<1e^−4^) (**Figure S9-S10**). We estimate the rate, through linear regression, over the global alignment to 25.1 (23.3 – 27.2) substitutions per genome per year, with significantly slower estimates of 18.3 (17.3 – 18.9) for the sub-sampled B.1.1 group suggesting some slowing down of rates through time since the beginning of the pandemic^23,24^. To formally estimate the age of common ancestor of the B.1.1.50 clade, dominated by samples from our study, we applied TreeDater^25^ to the subsampled B.1.1 phylogeny (**Figure S11**). This allowed us to estimate the age of the node giving rise to the B.1.1.50 grouping to the 5^th^ of February 2020 (16^th^ January – 19^st^ February; confidence intervals following parametric bootstrapping) (**Figure 2**). This suggests that Palestinian strains belonging to this clade were being transmitted locally already around these dates. Our timed phylogeny further allows estimation of the lower bound of the date of introduction into various local regions. For example, the collection of samples from the Nablus governorate share an estimated most recent common ancestor dating to the 24^th^ of May (13^th^ of May – 2nd of June) and the three genomes isolated from patients on the 22nd of June in Halhoul share an estimated ancestor dating to the 9^th^ of June (28^th^ May – 17^th^ June).

**Figure 2.**
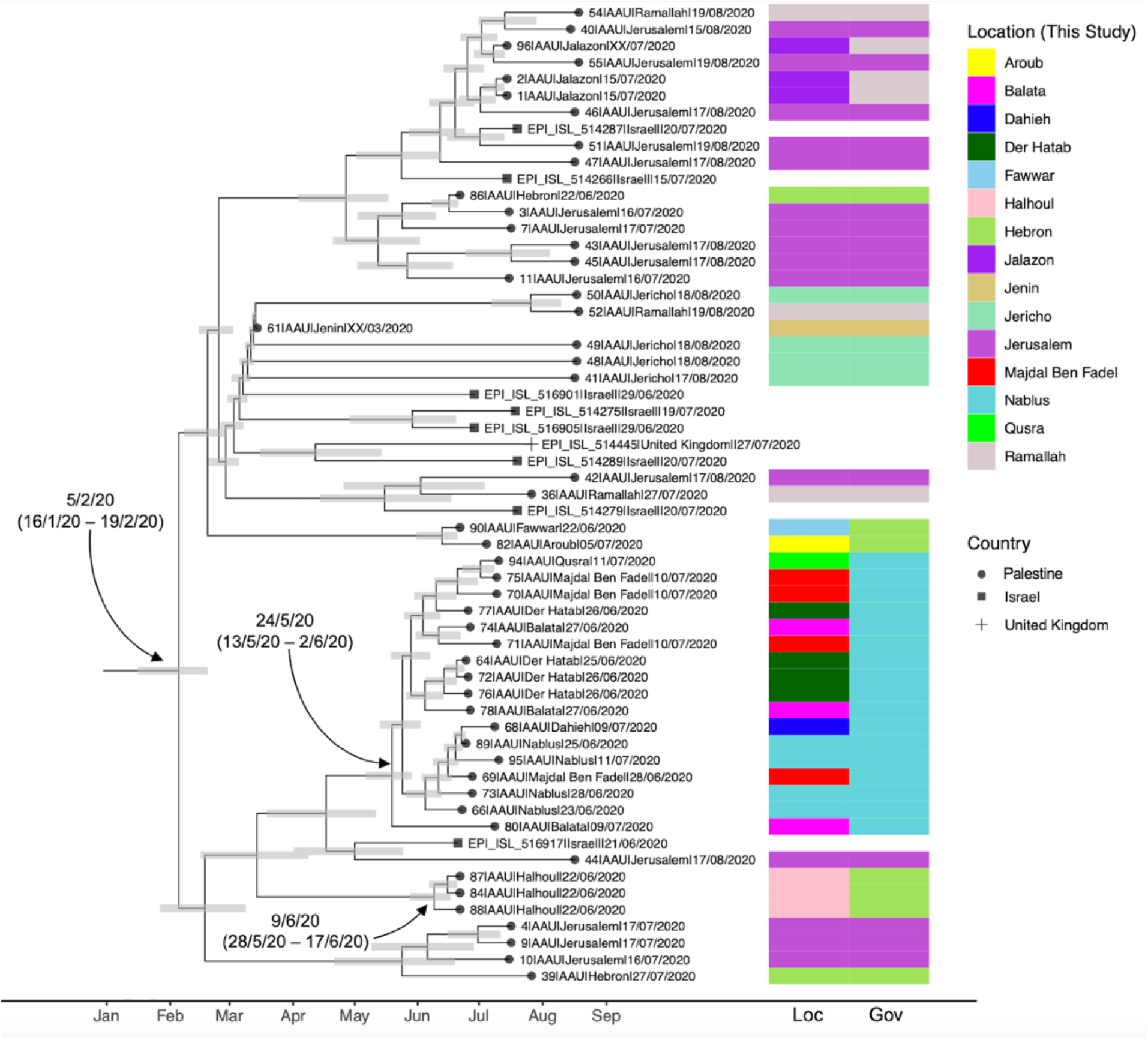
Time calibrated phylogenetic tree for the closely related B.1.1.50 clade subset from 1,252 B.1.1 genomes. The coloured panel at right provides the location (‘Loc’) and Governorate (‘Gov’) of samples generated in this study (Palestine). Samples without colour panels derive from Israel and the United Kingdom as given in the tip labels. Grey bars provide the 95% confidence intervals around the estimated age of phylogenetic nodes.

### Intra-isolate minor allelic diversity

After discarding minor alleles with an intra-individual frequency <0.05, which for most are likely to be spurious, 598 polymorphic sites were retained. The vast majority of minor alleles (96%) displayed frequencies 0.05<x<0.2 (**Figure S12 and S13**) and most of them were present only in one sample (**Figure S14**). The low frequency of minor alleles shared between samples prevented us from using this signal to reconstruct transmission chains. Indeed, we found no statistically significant correlation between the SNP-based phylogenetic signal and that of minor alleles (R2 = 7.7e^−4^; permutation *p*-value=0.94). The lack of congruence between SNP-based phylogenetic signal and the distribution of minor allele frequency variants suggests that it would be difficult to leverage the latter for the reconstruction of transmission chains (**Figure S15** and **Figure S16**).

## Discussion

Phylogenetic analyses of a SARS-CoV-2 genomic dataset from Palestine point to an earlier introduction and circulation of the virus than had been previously recognised, in line with the situation in many other regions of the world. This suggests SARS-CoV-2 was in sustained circulation prior to the establishment of public health interventions. The local COVID-19 epidemic(s) in Palestine were seeded by multiple (at least nine) independent introductions of SARS-CoV-2, though the lack of geographic structure and incomplete sampling make it challenging to pinpoint the exact sources of import and export events. However, the overarching diversity in circulation reflects that SARS-CoV-2 in Palestine recapitulates at least some of the global diversity in the SARS-CoV-2 population, though we do identify cases of more local community transmission.

One of the key challenges in reconstructing the spread, and more formal direct transmissions, of SARS-CoV-2 is its relatively low mutation rate, meaning multiple transmissions can occur before any mutation occured^26^. Our estimated rate over the global alignment of 25.1 (23.3-27.2) mutations per genome per year falls in line with other published rates, and remains consistent with mutation rates observed in other coronaviruses which are maintained relatively low due to the action of a proof-reading protein (non-structural polyprotein 14)^27^. Epidemiological reconstructions are further challenged by the rapid global dissemination of SARS-CoV-2 and its unbalanced geographical sampling. Care must therefore be taken when assigning geographic origins to imported cases^17,26^. As an example, our dataset includes three samples falling into a B.1 clade of 110 genetically identical sequences sampled over 21 nations over the course of 53 days.

A possible approach to reconstructing transmission in these settings has been suggested by the use of shared minority variants^18,28^. In our dataset we do identify a set of three identical isolates, two of which share minority variants, suggesting these two samples are more closely related. However, overall, despite considerably deep sequencing of the samples in our dataset, we find no usable phylogenetic signal in minor allelic variants that may be leveraged to aid in the reconstruction of transmission chains. Indeed, we found no evidence for any correlation between pairwise genetic distance between samples and their propensity to share minority variants.

The bulk of SARS-CoV-2 genomes in our dataset fell into a single cluster of B.1.1.50 SARS-CoV-2. Not precluding the possibility of many unsampled cases, our phylogenetic analyses points to this phylogenetic grouping representing a major local transmission cluster which has accumulated diversity primarily within Palestine. This cluster includes one SARS-CoV-2 sampled from the UK and eight from patients in Israel; with 18,139 and 288 representatives from these countries in our global dataset respectively.

We estimate the age of this cluster to early February, predating the first reports of COVID-19 positive patients in Palestine in a hotel in Bethlehem, where a group of Greek tourists had visited the hotel in late February and were diagnosed with the virus, as well as confirmed cases of college students returning from Europe. The cluster has been maintained until at least the latter half of August of 2020. Consequently, other local phylogenetic clusters have emerged from within the initial B.1.1.50 clade, for example the sampled viral sub-population circulating in the Governorate of Nablus since mid to late May.

Due to the limited geographic structuring of the global genetic distribution of SARS-CoV-2, it is difficult to confidently identify putative sources of introduction of the virus to particular locally transmitting lineages. Thus, we can only speculate on the origin of the possible introductions. On the basis of human movement, Israel and Europe represent the most plausible sources. Despite attempts by the Palestinian government to discourage its residents from crossing from and into Israeli areas, daily commuting of workers and residents between the West Bank and Israel never entirely ceased with. Also, close to 2000 Palestinians entered the West Bank from Jordan via the Allenby crossing between the 1st and 13th of July. Another plausible source are Palestinian students returning from Europe as well as the USA.

At the same time, our genomic analysis pinpoints the presence of transmission lineages for which there are no known epidemiological links, for example we identify a clear case of localised community transmission predating by at least weeks the earliest cases in Palestine. As such, our study supports the adoption of genomic surveillance in Palestine, highlighting the potential of genomic epidemiology to uncover and ultimately monitor patterns of disease spread at both global and local scales.

## Materials and Methods

### Data Collection and Processing

Nasopharyngeal swabs were sampled between the 4th of March and the 19^th^ of August 2020 from 69 symptomatic and asymptomatic Palestinian patients originating from 17 locations within eight governorates (**Table S1**, **Figure S1**). RNA was extracted from clinical samples using using QIAamp MinElute virus spin kit (Catalogue number 57704). Real-time reverse transcriptase (RT)-PCR was used to detect SARS-CoV-2 with Ct values ranging between 9-30. All specimens were handled under a biosafety cabinet according to laboratory biosafety guidelines. For four samples (60, 61, 62, 96), for which only information on the month of sample collection was available, the collection date was set to the middle (15^th^) of the month. Approval from National ethical committee was obtained (PHRC/HC/738/20).

### Palestine SARS-CoV-2 dataset: sequencing and variant calling

cDNA synthesis was done using NEBNext^®^ non-directional RNA-Seq workflow and NEBNext Ultra RNA First Strand Synthesis Module ([NEB #E7771](https://international.neb.com/products/e7771-nebnext-ultra-ii-rna-first-strand-synthesis-module)) and NEBNext^®^ RNA Second Strand Synthesis Module ([NEB #E6111](https://international.neb.com/products/e6111-nebnext-ultra-ii-non-directional-rna-second-strand-synthesis-module)). Library preparation was performed with the Nextera Flex for Enrichment workflow^29^. Sequencing was performed on Illumina Next-Seq 550 sequencing system. The mapping of the raw sequencing reads to the Wuhan-Hu-1 reference sequence (GenBank accession MN908947; equivalent GISAID ID EPI_ISL_402125) was performed using the Dragen RNA pathogen detection v.3.5.14 pipeline^30^. Strains displayed a mean coverage ranging from 22 to 9400X (**Table S1**). Variants were then called using Freebayes v1.22^31^. Three distinct indels (each present in less than three isolates) and 762 unfiltered variable sites were identified. We called SNPs supported by an intra-isolate frequency of 0.65 for further phylogenetic study, but we lowered this threshold to 0.05 to create a SNP set dedicated to the study of minor allelic frequencies. SNPs flagged as putative sequencing errors were masked (a full list of ‘masked’ sites is available at https://github.com/W-L/ProblematicSites_SARS-CoV2/blob/master/problematic_sites_sarsCov2.vcf, accessed 25/08/2020)^32^. Across the final dataset we obtained 128 high-quality SNPs.

### Worldwide SARS-CoV-2 dataset

Additionally, we downloaded 56,803 high quality assemblies (high coverage, >29,700bp and with a fraction of ‘N’ nucleotides <5%.) from the worldwide SARS-CoV-2 diversity available on GISAID^4,5^ on 25/08/2020. All animal isolate strains were removed as well as samples flagged by NextStrain as ‘exclude’ (https://github.com/nextstrain/ncov/blob/master/config/exclude.txt as of 25/08/2020). This left 54,793 assemblies for downstream analysis. A full metadata table, list of acknowledgements and exclusions is provided in **Table S3**. The 54,793 SARS-CoV-2 assemblies were profile aligned against Wuhan-Hu-1 (MN908947.3) using MAFFT v.7.205^33^.

### Phylogenetic reconstruction

The 69 aligned Palestinian sequences generated herein and 54,793 strains from the worldwide diversity were concatenated and a maximum likelihood tree built using IQ-TREE 2.1.0 Covid release^34^. A further 57 long-branch phylogenetic outliers were removed following application of TreeShrink^35^ (given in **Table S3**). The final tree of 54,804 samples, rooted on Wuhan-Hu-1, is provided in **Figure S3**. Trees were queried and plotted using the R packages Ape v5.4 ^36^ and ggtree v1.16.6^37^.

### Lineage assignment and mutation analysis

Lineages were assigned to each of the Palestinian SARS-CoV-2 assemblies using the dynamic nomenclature tool Pangolin^22^ (https://github.com/cov-lineages/pangolin, applied 28/8/2020). The nucleotide position of SNPs identified in the 69 assemblies is provided in **Table S2**, with annotations provided relative to Wuhan-Hu-1. 67/69 assemblies carried the derived D614G mutation in the spike protein, with 65 carrying the full four mutation D614G haplotype (nucleotide positions 241, 3037, 14408, 23403). 52 also carried the three neighbouring mutations in the nucleocapsid protein (28881-28883). The number of SNPs differing between viral assemblies within Palestine and across global datasets was assessed using SNP-sites^38^ and SNP-dists (https://github.com/tseemann/snp-dists) with heatmaps plotted using ComplexHeatMap v2.1.2^39^.

### Phylogenetic Dating

To estimate the age of the largest transmission cluster we extracted a subset of 1,252 B.1.1 SARS-CoV-2 from the phylogeny including the B.1.1.50 clade. The BactDating^40^ *roottotip()* function was applied to compute the root-to-tip temporal regression for both the global tree and subsets of trees. In all cases significance was assessed following 10,000 random permutations of sampling dates. Confidence intervals around the inferred rates were assessed through 1,000 bootstrap resamples with replacement. As with the global tree, the subset B.1.1. clade exhibited measurable evolution through time both with and without the earliest SARS-CoV-2 genome (reference Wuhan-Hu-1) included (p<1e^−4^ in all cases). Following confirmation of significant temporal signal we applied *dater()* within the TreeDater package v0.50 specifying a strict clock model and assessed confidence intervals following 100 iterations of the *parboot()* parametric bootstrap fitting method. Tip-dated phylogenetic trees, together with associated confidence intervals, were assessed and plotted using ggtree v1.16.6^37^.

## Data Availability

All newly generated assemblies have been uploaded to GISAID and are available under IDs: EPI_ISL_596500 - EPI_ISL_596568. In addition, raw short reads have been uploaded to the NCBI Short Read Archive (SRA) under BioProject PRJNA669945.

## Acknowledgements and Funding

N.Q., Z.S., H.D. and H.S. acknowledge project financial support from AAUP. L.v.D and F.B. acknowledge financial support from the Newton Fund UK-China NSFC initiative (grant MR/P007597/1) and the BBSRC (equipment grant BB/R01356X/1). L.v.D is supported by a UCL Excellence Fellowship. DR is supported by a NIHR Precision AMR award. We would like to thank the Molecular Genetics and Gene Toxicology Masters students at AAUP, Iman Bouarfa, Rawan Obeid and Niveen Bdier for discussions. We additionally wish to acknowledge the large number of originating and submitting laboratories who have readily shared genome assemblies on GISAID. A full table of acknowledgements can be found in **Table S3**.

