## Supplementary figures for "The genomic epidemiology of SARS-CoV-2 in Palestine"

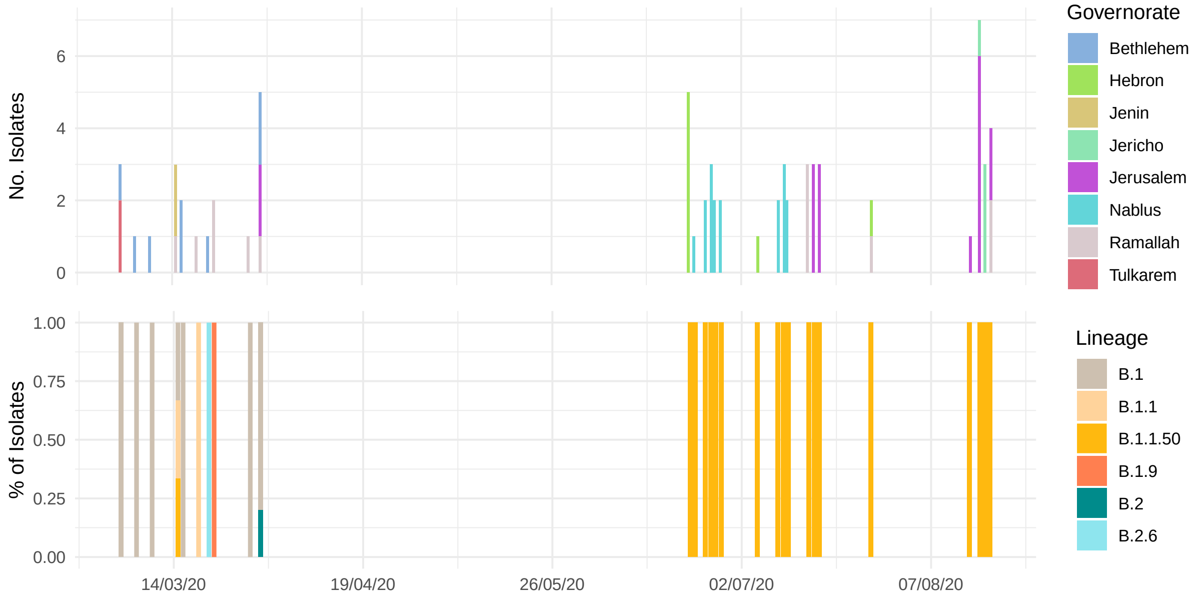


**Figure S1** (top) Number of SARS-CoV-2 isolates obtained over time (x-axis) coloured by the region Governorate. (bottom) Pangolin lineage assignments (%) obtained for each sample. Full metadata for all novel data presented in this study is provided in **Table S1**.

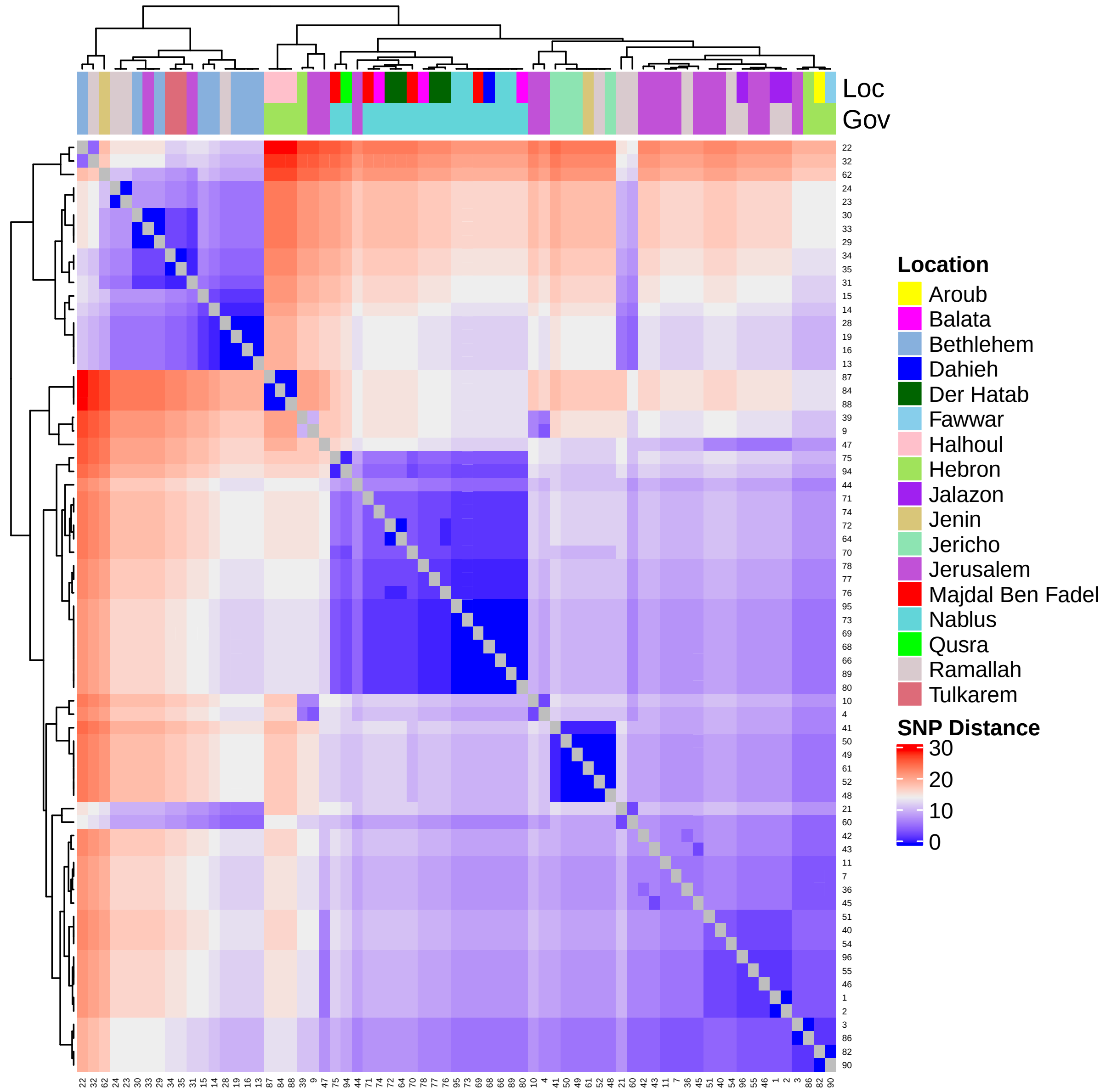


**Figure S2** Pairwise SNP differences between all SARS-CoV-2 data collected in this study. Colour provides the number of SNPs differing as per the legend at right. The top panel provides the location and governorate where samples were collected.

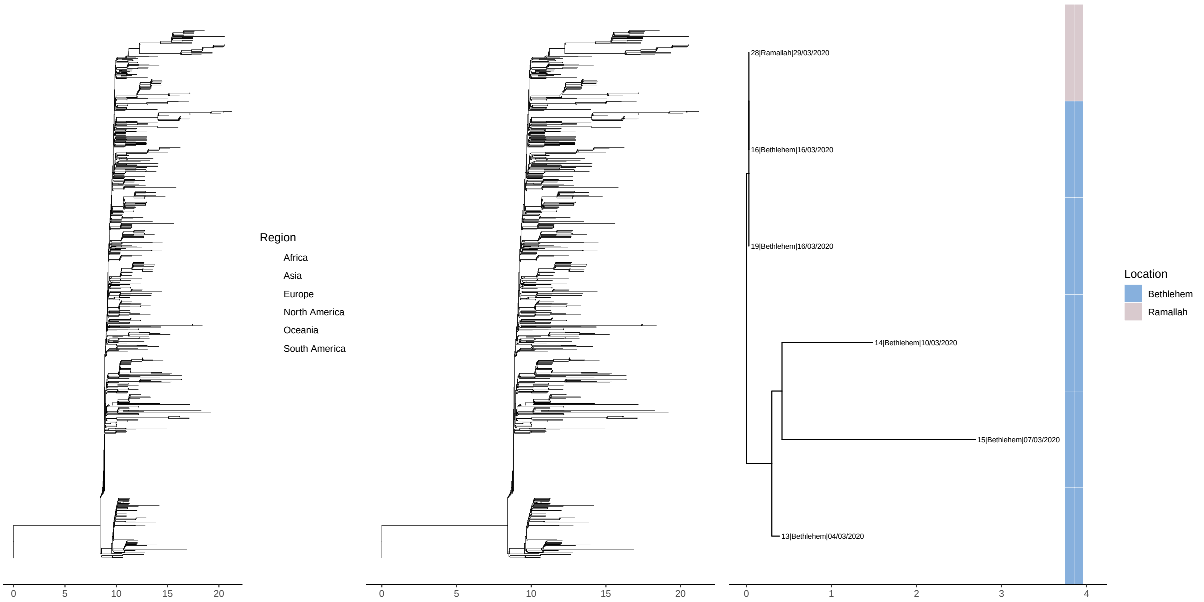


**Figure S3** B.1 node extracted from the global SARS-CoV-2 phylogenetic tree including six samples from this study. The tree at left provides the global phylogenetic context with samples from this study highlighted in the middle panel. The panel at right provides a subset of the tree to demonstrate the diversity amongst our samples with the coloured bar providing the location/governorate.

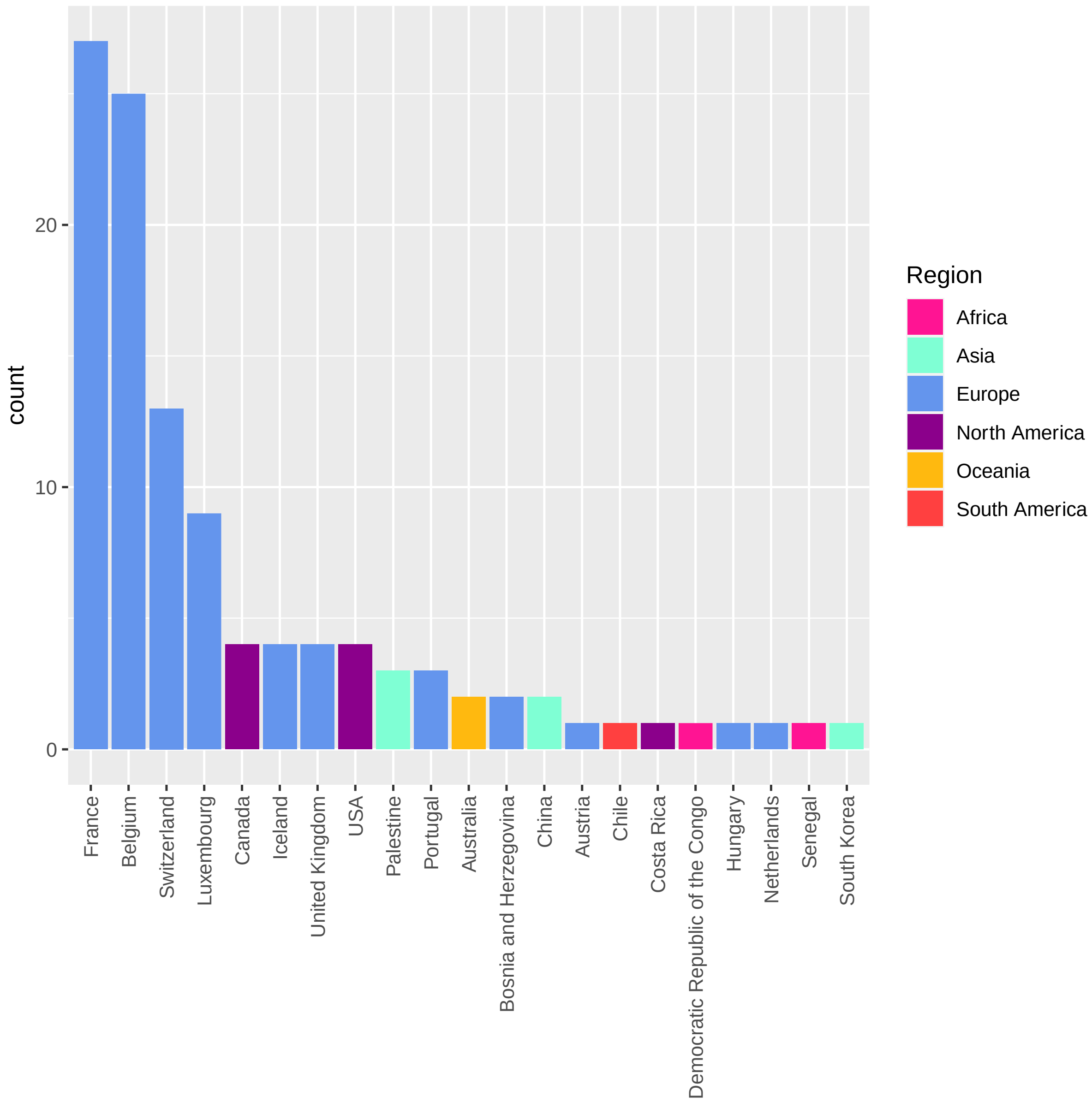


**Figure S4** Geographic origin of the of the110 SARS-CoV-2 strains with identical genome sequences (B.1 lineage) including three Palestinian samples.

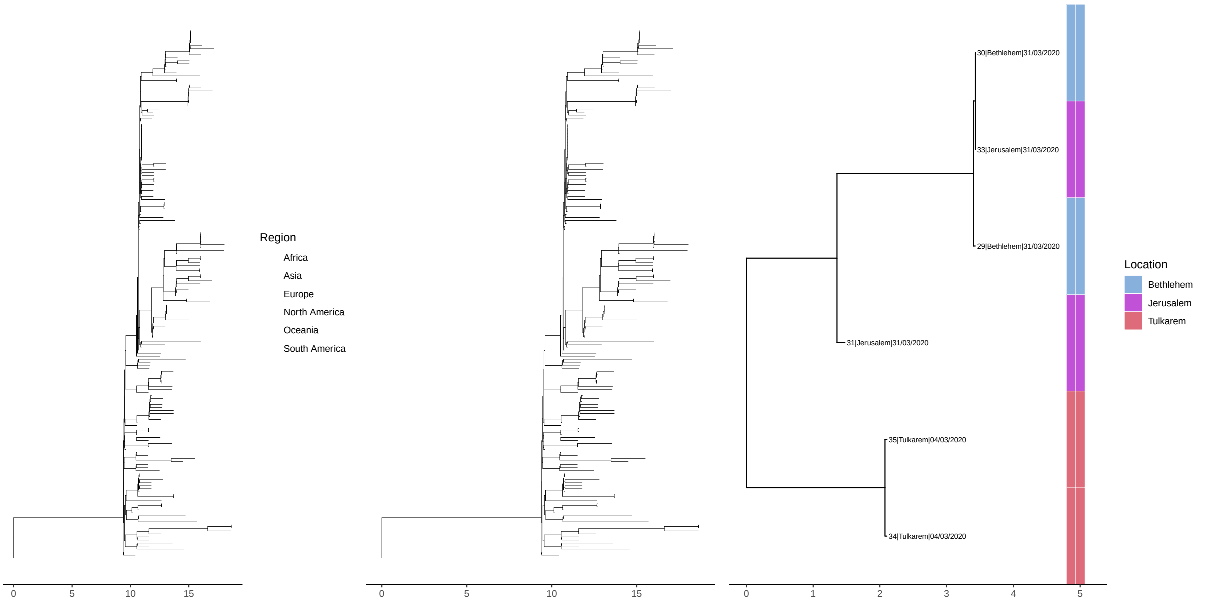


**Figure S5** B.1 node extracted from the global SARS-CoV-2 phylogenetic tree including six samples from this study. The tree at left provides the global phylogenetic context with samples from this study highlighted in the middle panel. The panel at right provides a subset of the tree to demonstrate the diversity amongst our samples with the coloured bar providing the location/governorate.

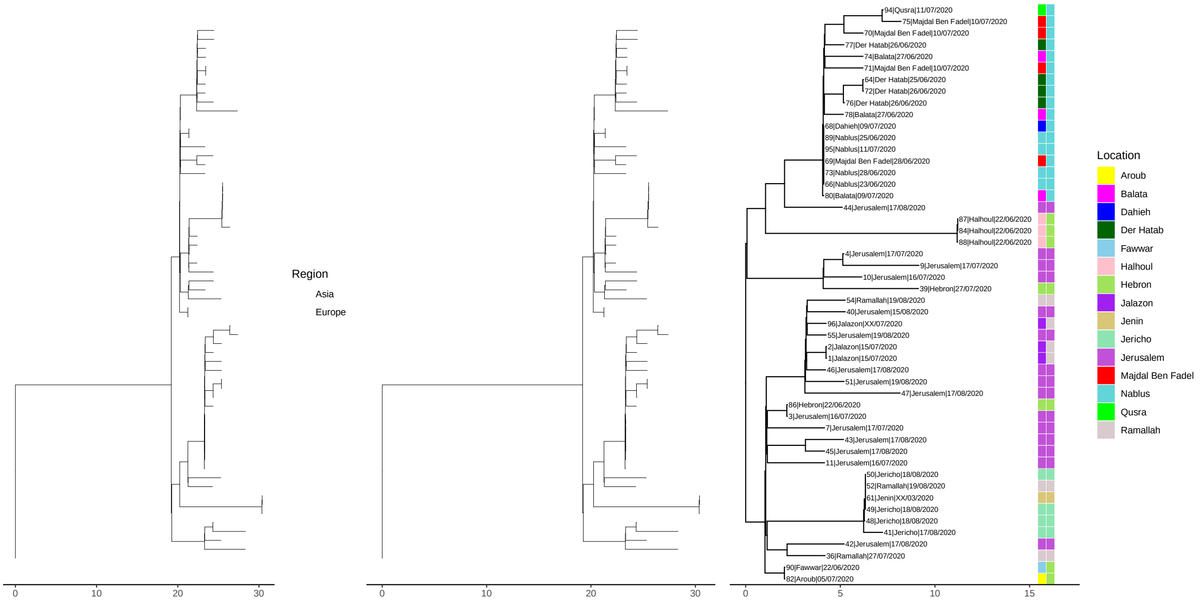


**Figure S6** B.1.1.50 node extracted from the global SARS-CoV-2 phylogenetic tree including fifty samples from this study. The tree at left provides the global phylogenetic context with samples from this study highlighted in the middle panel. The panel at right provides a subset of the tree to demonstrate the diversity amongst our samples with the coloured bar providing the location/governorate.

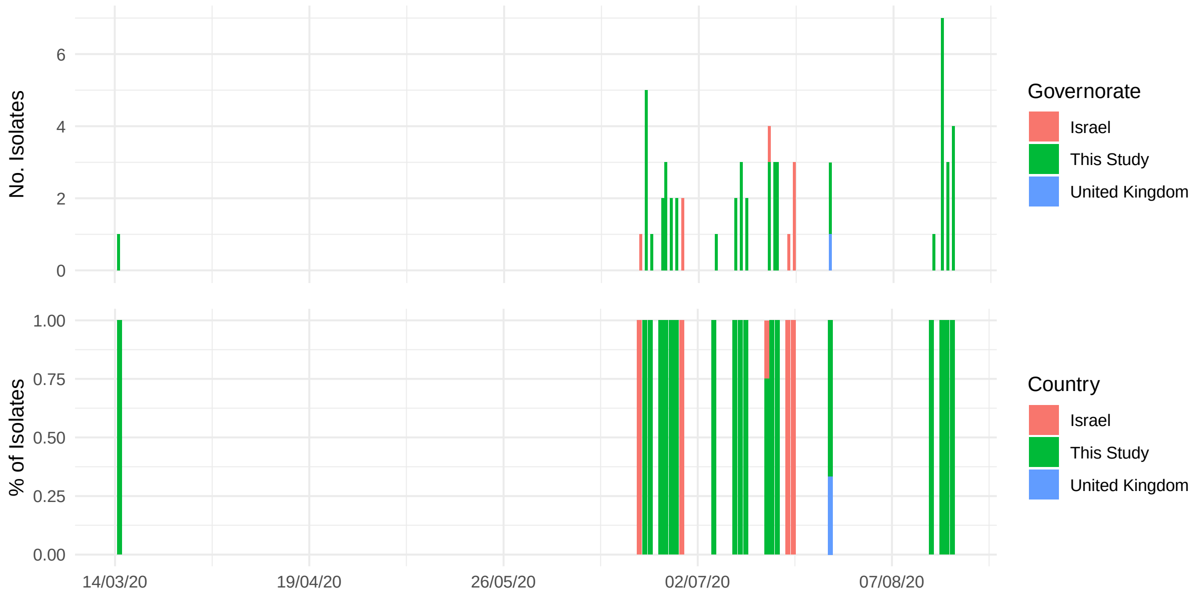


**Figure S7** Sample collection dates and nation of isolation for SARS-CoV-2 isolates falling within the B.1.1.50 local phylogenetic cluster (**Figure S6**).

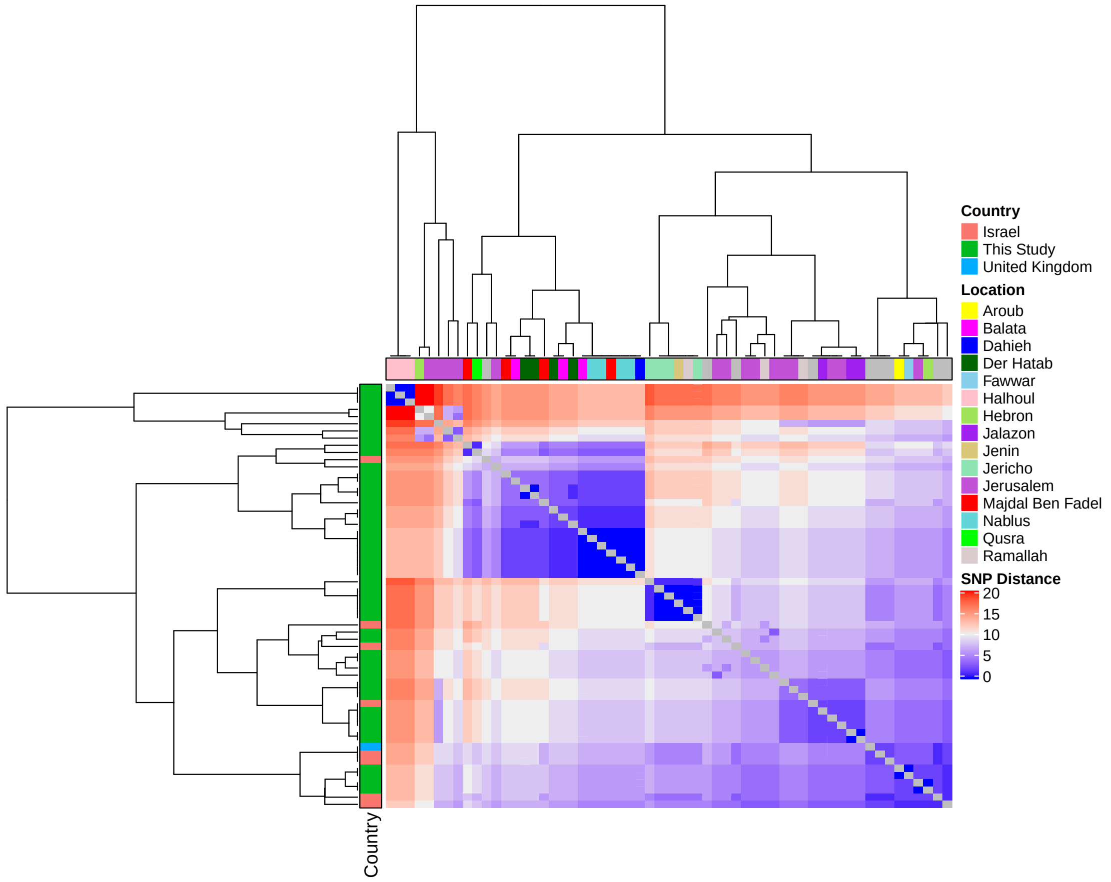


**Figure S8** Pairwise SNP differences between SARS-CoV-2 falling within the B1.1.50 local cluster, including 50 SARS-CoV-2 genomes generated in this study. The colour scale provides the number of SNPs differing as per the legend at right. The top panel provides the location within Palestine (samples not generated in this study set to grey) where samples were collected. The panel at left provides the country assignment (nine global samples from Israel [n=8] and the UK [n=1]).

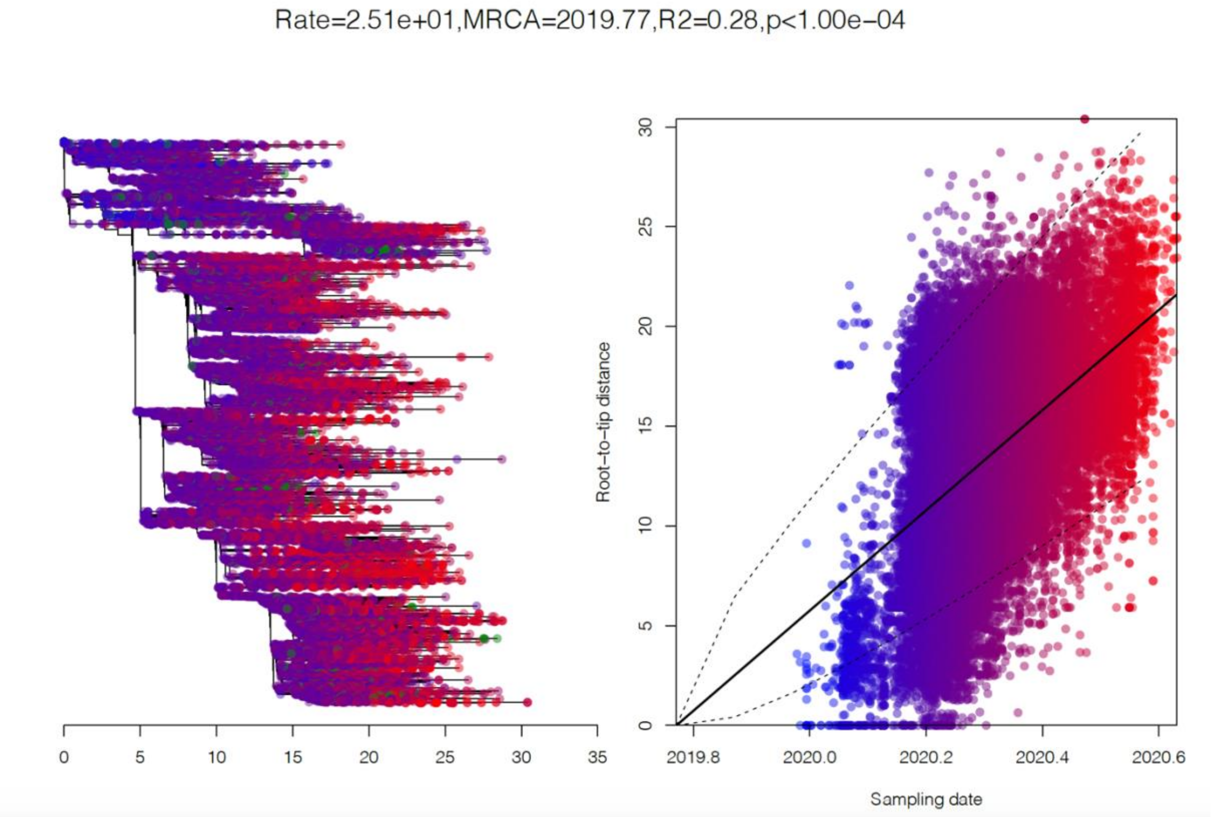


**Figure S9** (left) Global phylogenetic tree rooted on Wuhan-Hu-1. (right) Correlation between the decimal date of sample collection (x-axis) and root-to-tip phylogenetic distance (y-axis) across the global dataset, with points coloured by location of collection. A significant temporal regression was obtained following 10,000 randomisations of collection date (MRCA 2019.77, $r^2$=0.276, p<1e-4). Estimated rates following 1000 bootstrap resamplings gave rise to values of 25.1 (23.3 – 27.2) subsitutions per genome per year.

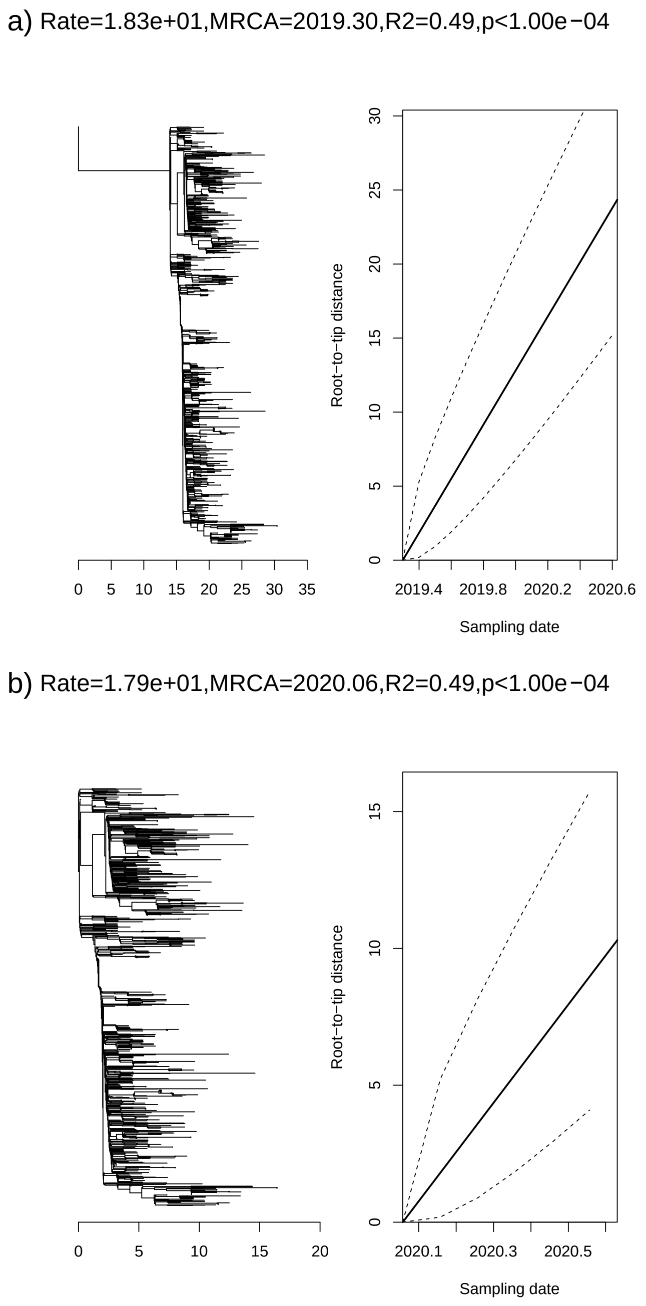


**Figure S10:** Subset phylogenetic tree of 1,252 B.1.1 isolates. a) Left panel provides the phylogenetic tree rooted on Wuhan-Hu-1. The right panel provides the root-to-tip phylogenetic distance (y) as a produce of the time of sampling (x), displaying a significant regression of rate 18.3 mutations per genome per year. b) Left panel provides the phylogenetic tree now dropping Wuhan-Hu-1. The right panel provides the root-to-tip phylogenetic distance (y) as a produce of the time of sampling (x), displaying a significant regression of rate 17.9 mutations per genome per year. In both cases *p*-values are provided following testing against 10,000 randomisations of the sampling date.

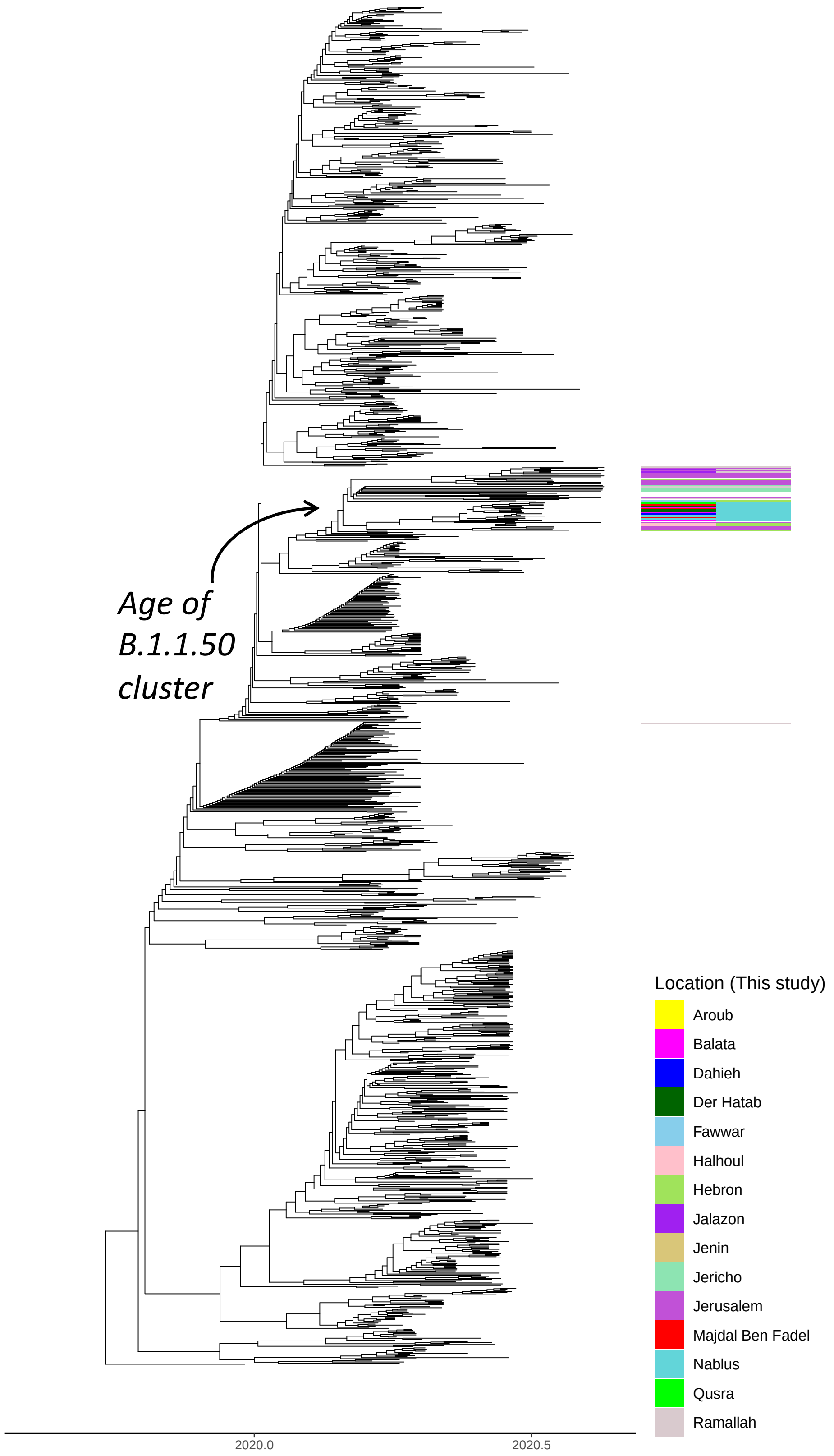


**Figure S11:** TreeDater time calibrated phylogenetic tree of 1252 B.1.1 SARS-CoV-2 isolates. Data from this study (Palestine) are highlighted by the coloured bars which give the location and governorate of patient location. Grey bars provide the 95% confidence intervals following parametric bootstrapping. The highlighted age of the B.1.1.50 clade dominated by data from our study is 5$^{th}$ February (16th January – 19th of February) (main text **Figure 2**).

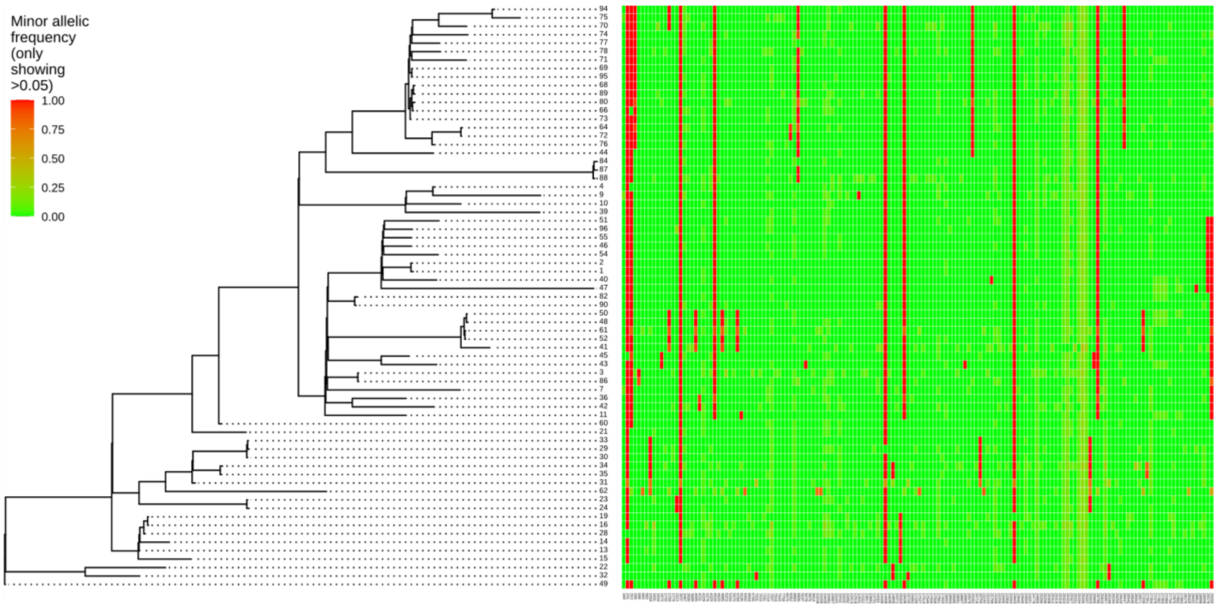


**Figure S12:** Alignment of the 69 genomes from Palestine restricted to the 167 sites with derived allele at frequency >0.05 in at least two isolates. The frequency of the minor variants is shown as a heat map. the vast majority of minor alleles (96%) displayed frequencies 0.05<x<0.2. Derived alleles near or at fixation are displayed in red.

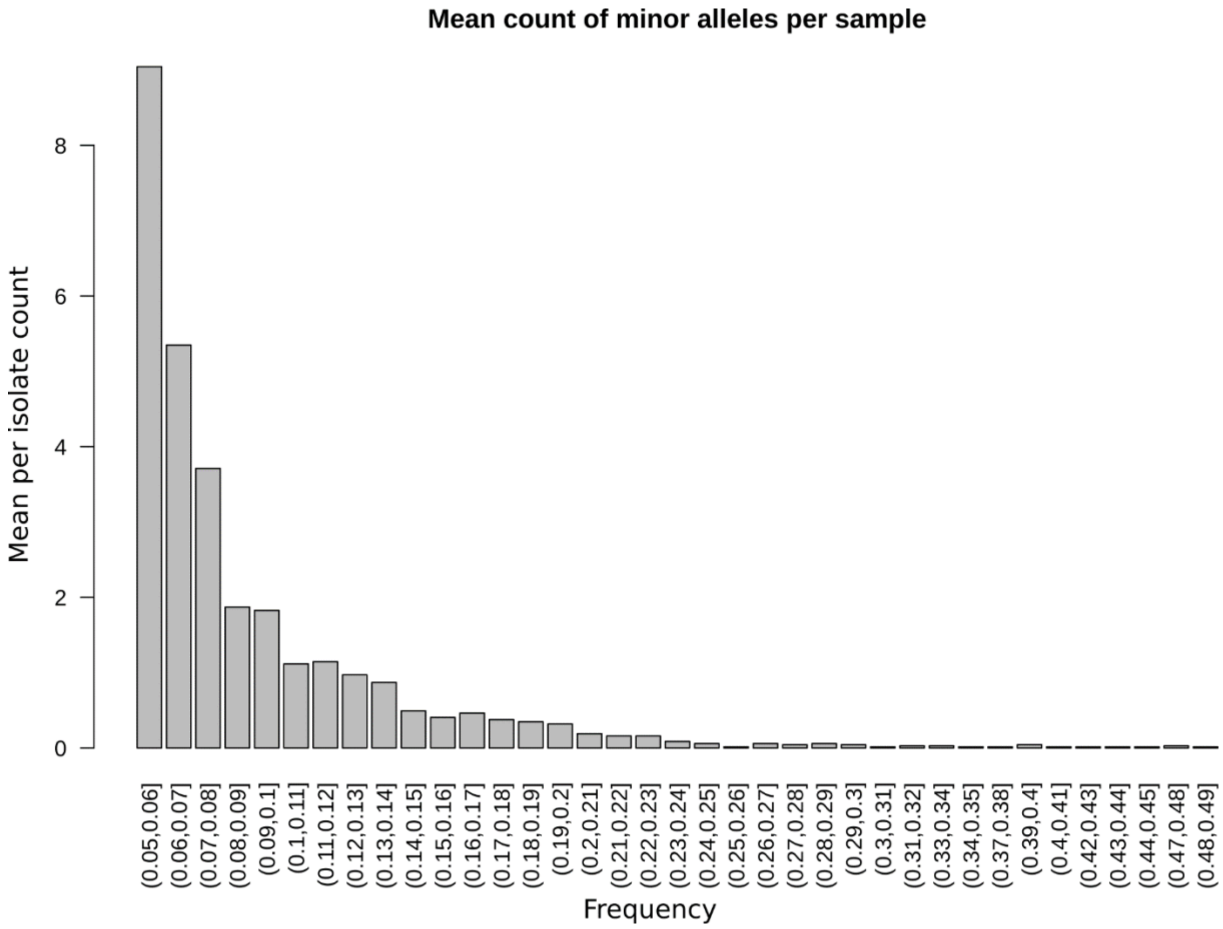


**Figure S13:** Distribution of the frequency of minor allele variants of within host polymorphisms found at a frequency above 0.05.

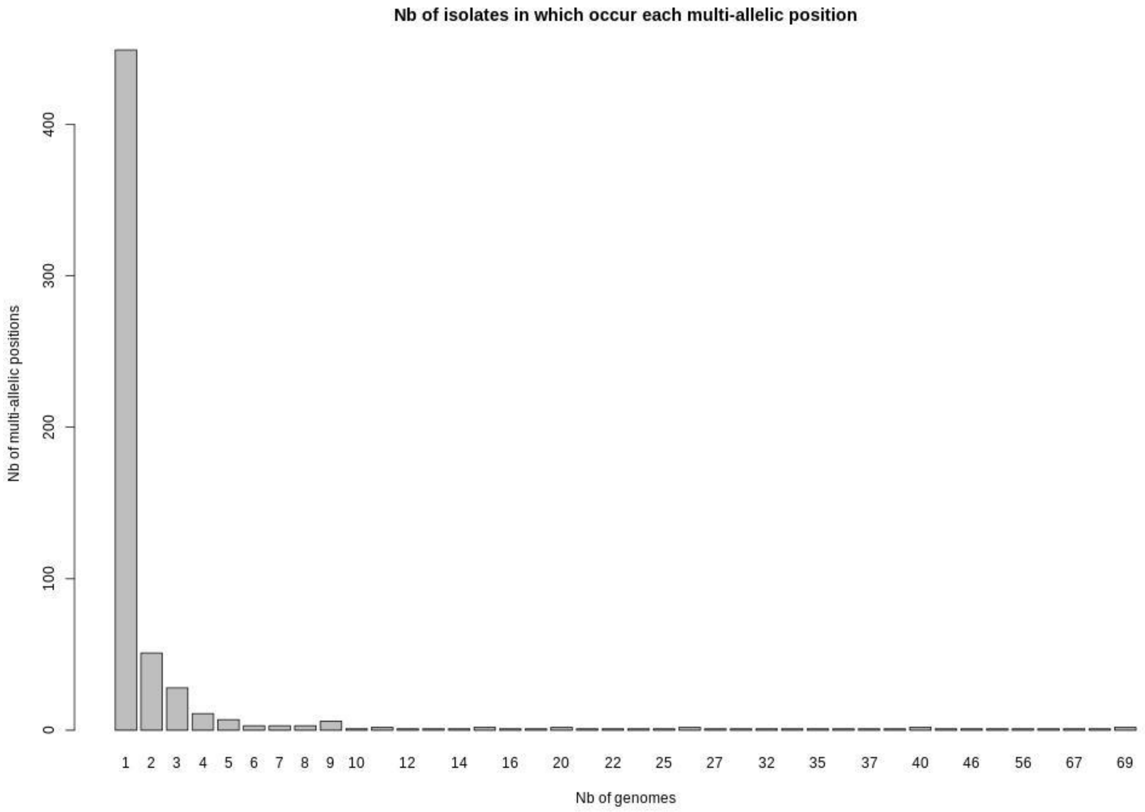


**Figure S14:** Distribution of the number of SARS-CoV-2 genomes sharing individual minor allele variants at a frequency above 0.05.

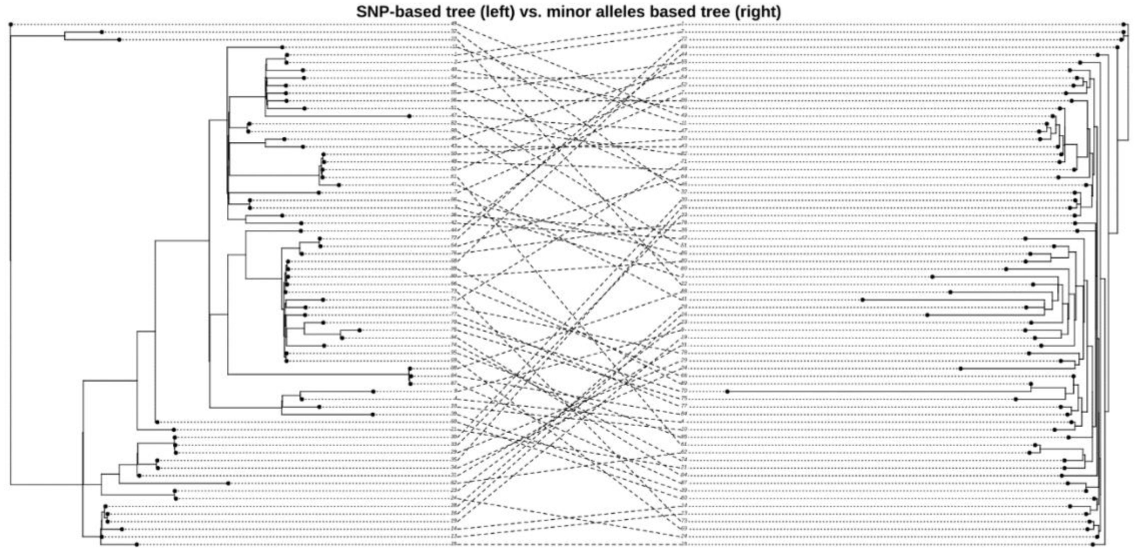


**Figure S15:** Cophyloplot showing the correspondence of the position of the strains in a maximum-likelihood SNP-based tree (left) and a maximum-likelihood minor allele-based tree (right). The minor allele-based tree is based on the binary matrix of presence or absence of a minor allele (frequency >0.05) at each nucleotide position of the genome.

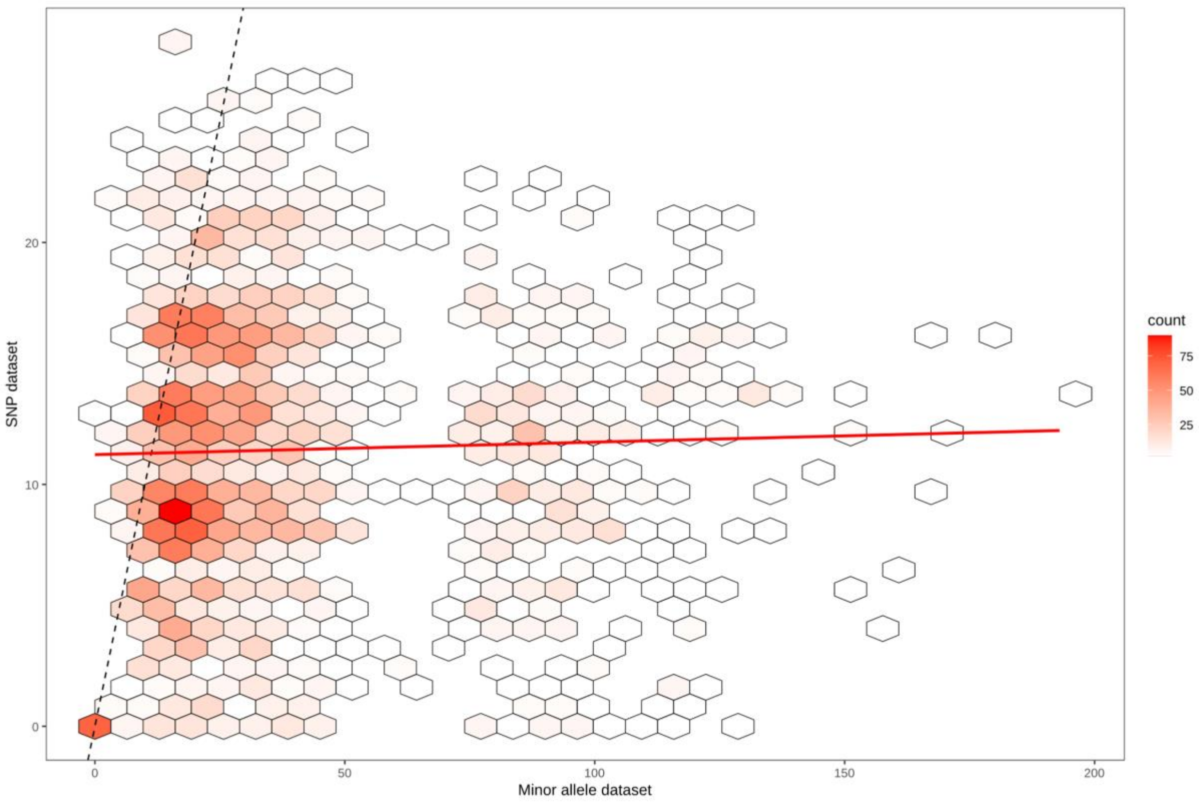


**Figure S16;** Correlation between the pairwise distances between SARS-CoV-2 genomes in the SNP-based alignment and the minor-allele based alignment matrices. Red line : linear regression ; dashed line : y=x. The relation between SNP-based and minor-allele paiwise genetic distances was not statistically signioficant (R2 = 7.7e$^{-4}$; Mantel test *p*-value=0.94).
